# Simple theoretical models predict some - but not all – aspects of the experimental evolution of antibiotic resistance

**DOI:** 10.1101/2025.09.07.674561

**Authors:** Elin Lilja, Rosalind J. Allen, Bartlomiej Waclaw

## Abstract

Mathematical modelling of antibiotic resistance plays an important role in understanding the mechanisms of resistance emergence and spreading, testing the feasibility of new treatment protocols, and antimicrobial stewardship. However, many assumptions underlying some of the most commonly used mathematical models have not been rigorously tested experimentally. We verify whether one of these models - a birth-death-mutation process - is able to quantitatively predict the outcome of laboratory experiments. We grow bacteria in a bioreactor in conditions that closely resemble the assumptions of the model, and compare the model predictions with experimental observables such as the probability and time to resistance evolution, mutant number distribution, and the genetic composition of the evolved populations. We show that the model fails to reproduce some aspects of the experiments (failing differently for different antibiotics) but that simple modifications of the model significantly improve its predictive power. These modifications give insight into the population dynamics of resistant mutants for each antibiotic tested, and highlight the importance of quantitative modelling for accurate prediction of antibiotic resistance evolution.

## Introduction

Evolution of bacterial resistance to antibiotics is universally recognized as one of the most urgent problems in medicine [1–3]. Both in-vitro laboratory models of resistance evolution [4–9] and mathematical models [10–17] are important tools for understanding the mechanisms of resistance emergence and spreading. *In-vitro* models enable the study of bacterial evolution in controlled conditions which cannot be achieved *in vivo*, while mathematical models can provide insight into processes that are difficult to observe directly. These approaches are most powerful when used in combination; however, while both *in vitro* and mathematical models have been used to qualitatively explain various aspects of antimicrobial resistance, quantitative comparisons between experiments and theory are rare [18–22]. This lack of quantitative validation may lead to misinterpretation of experimental data, since, for example, it may be hard to distinguish between mechanistic models that generate qualitatively similar outcomes. Lack of experimental validation is also a barrier to the evaluation of alternative theoretical models. More extensive quantitative comparison between *in-vitro* experimental data and mathematical models would greatly facilitate the development of predictive data-driven models for the emergence of bacterial antibiotic resistance.

The classic model that is often used to describe the dynamics of bacterial antibiotic resistance evolution is the birth-death-mutation model [23,24]. This model considers sensitive and resistant subpopulations of cells which grow and die; the growth and death rates may differ between the resistant and sensitive strains and in the presence or absence of antibiotic. Furthermore, sensitive cells mutate into resistant cells with a small probability. The simple model has been extended in many ways, including non-exponential growth [25] and spatial selection gradients [26,27]. Example applications include the evolution of resistance during long-term antibiotic treatment [28], mutational pathways to resistance [29], and strain competition during combination therapy [30].

In this work we ask to what extent the birth-death-mutation mathematical model can quantitatively reproduce the evolution of antibiotic resistance in a bioreactor model of antibiotic treatment. We focus on the first step in the evolution of antibiotic resistance: the emergence and fixation of a single-point mutation that confers antibiotic resistance. We use a laboratory strain of *Escherichia coli* (the K-12 strain MG1655) and three different antibiotics: rifampicin (RIF), ciprofloxacin (CIP), and streptomycin (STR), each of which has a different mode of action. We designed our experiment to be as simple and as reproducible as possible, in order to facilitate mathematical modelling. Specifically, the population of bacteria grows exponentially, non-growing bacteria are diluted out, nutrients are unlimited, and the antibiotic concentration is at least 4x the minimum inhibitory concentration (MIC) to ensure that only resistant bacteria are able to proliferate in the presence of the antibiotic. This should eliminate mechanisms such as persistence or non-genetic resistance that could confound the results.

We show that the archetypal model of resistance evolution – the birth-death process with mutations – is able to reproduce some, but not all, aspects of the evolutionary dynamics of *de novo* resistant variants in our experiments. Moreover, different antibiotics violate different assumptions of the model. However, with the addition of simple modifications the model is able to quantitatively reproduce all of our experimental data. These modifications include: a delayed response to the antibiotic, multiple alleles having different growth rate, and phenotypic lag.

Our work confirms, reassuringly, that simple mathematical models can describe resistance evolution with quantitative accuracy. However, a quantitative description requires inclusion of the right biological mechanisms, which differ for different antibiotics. More broadly, our work shows how precise, quantitative measurements of bacterial growth combined with mathematical models can help to better understand evolutionary dynamics of bacteria and to construct better *in silico* models of AMR evolution.

## Results

### De novo evolution of resistance to rifampicin, ciprofloxacin and streptomycin in continuous cultures

We cultured *E.coli* cells in a bioreactor [31] in which bacteria, maintained in the exponential phase of growth by repeated cycles of dilution, were exposed to an antibiotic step-up (Fig. 1A). Our system is similar to the turbidostat [32] and morbidostat [8] but with the differences that the optical density is allowed to fluctuate and the concentration of the antibiotic, once added, is kept constant. To achieve steady exponential growth, the culture was diluted whenever the optical density exceeded 0.1, or every 30 minutes, whichever occurred first (Fig. 1B). The latter condition ensures that only actively growing bacteria are maintained in the culture; antibiotic sensitive cells, slowly-growing cells or persister cells are expected to be gradually diluted out upon antibiotic exposure. The bioreactor was inoculated with a mixture of two isogenic derivatives of the K12 *E. coli* strain MG1655: a YFP-fluorescent strain and a non-fluorescent strain, both with a deletion of the *fimA* gene to prevent biofilm formation in the culture bottles and in the cuvettes [31]. The use of a mixture of two easy-to-distinguish strains helped to determine the clonality of the final population.

**Figure 1.**
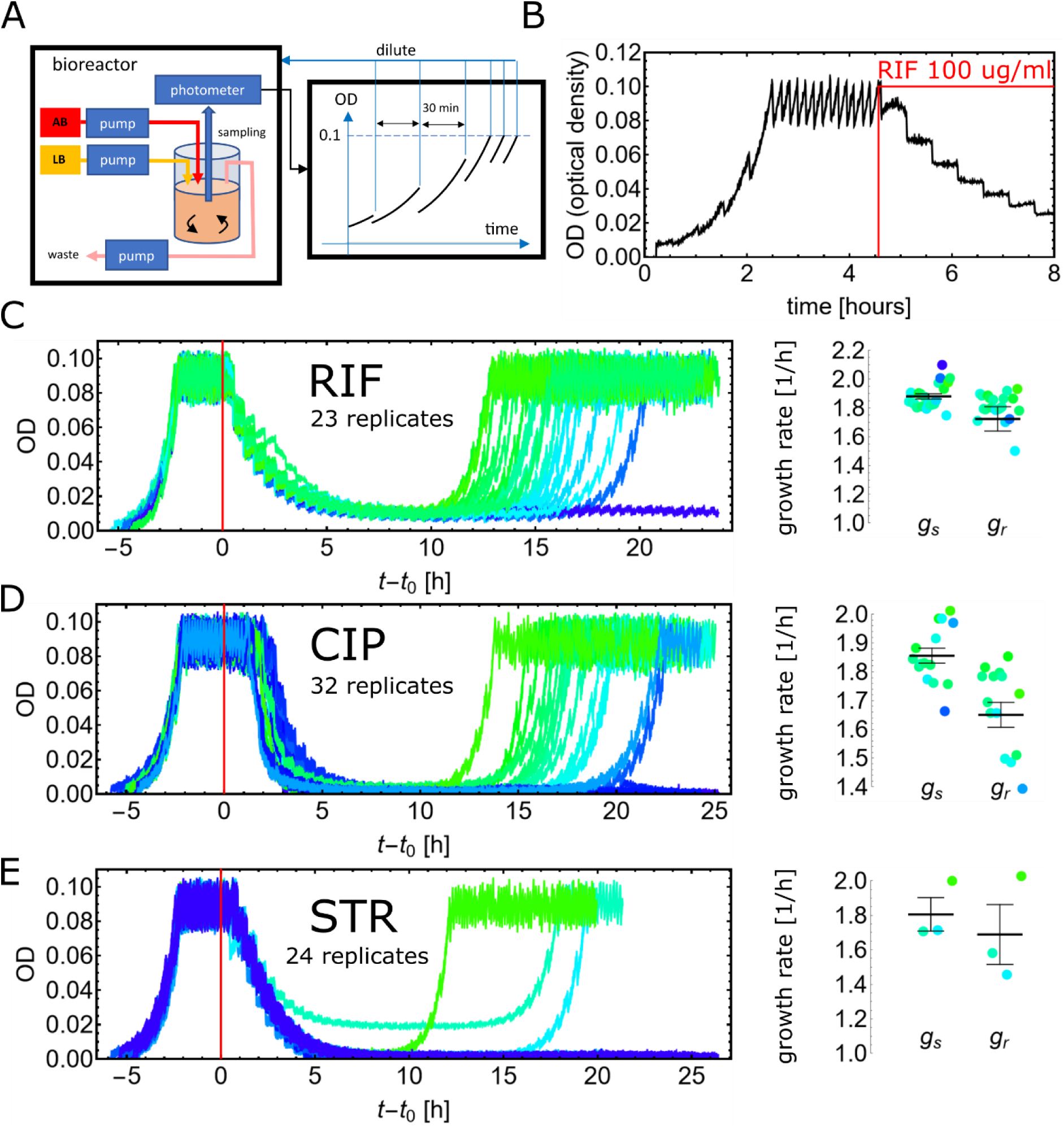
Evolution of resistance in a bioreactor. **(A)** Experimental setup. **(B)** The optical density over time in a typical experiment with rifampicin. The first part of the curve (*t* < 2.5 h) shows exponential growth with dilution every 30 min. The second part (2.5 < *t* < 4.5 h) shows rapid dilutions that keep the OD below 0.1. The last part (*t* > 4.5 h) shows the response to rifampicin. **(C-E)** Optical density versus time curves for all experiments, and the growth rates of sensitive and resistant populations obtained from these curves (Methods, “*Bioreactor data analysis*”), for rifampicin (C), ciprofloxacin (D) and streptomycin (E). The curves represent independent replicate experiments and have been colour-coded based on the regrowth time (green = fast regrowth, blue = slow or no regrowth). In one of the STR experiments, a problem with OD background correction caused the OD baseline to be visibly higher than zero.

We exposed bioreactor cultures of these antibiotic-sensitive strains to three different antibiotics: streptomycin, rifampicin and ciprofloxacin, in each case at concentrations several times above their MIC (specifically, we used 100 μg/ml streptomycin, 100 μg/ml rifampicin and 100 ng/ml ciprofloxacin). These antibiotics and concentrations were selected because, in each case, *E.coli* is known to be able to achieve the corresponding level of resistance via single point mutations in the target genes of the antibiotic, i.e., in *rpsL* (streptomycin) [9,33], *rpoB* (rifampicin) [34], and *gyrA* [35] (ciprofloxacin).

Figure 1C-E shows the optical density versus time for all our experiments, as well as the growth rates before and after antibiotic exposure. Regrowth after antibiotic exposure, indicating the emergence of resistance, occurred in 22/23, 15/32 and 3/24 of our rifampicin, ciprofloxacin and streptomycin experiments, respectively.

### A simple birth-death-mutation model cannot predict all experimental outcomes

To test whether a simple mathematical model could quantitatively explain our experimental data, we first considered the archetypal model for *de novo* evolution of antibiotic resistance: the birth-death process with mutations (Fig. 2A, inset). In our version of the model, we consider the dynamics of two subpopulations of bacteria that are respectively sensitive or resistant to the antibiotic. In the absence of the antibiotic, the sensitive cells are assumed to grow with rate *g*_S_(1 − *μ*) and mutate to become resistant to the antibiotic with rate *g*_S_*μ*, whereas the resistant cells grow with rate *g*_R_ that may be different to *g*_S_. Hence, in this model, mutation is coupled to growth. When the antibiotic is added to the continuously growing culture, all sensitive cells are assumed to immediately cease growth while resistant cells continue to grow with the same growth rate *g*_R_as before.

**Figure 2.**
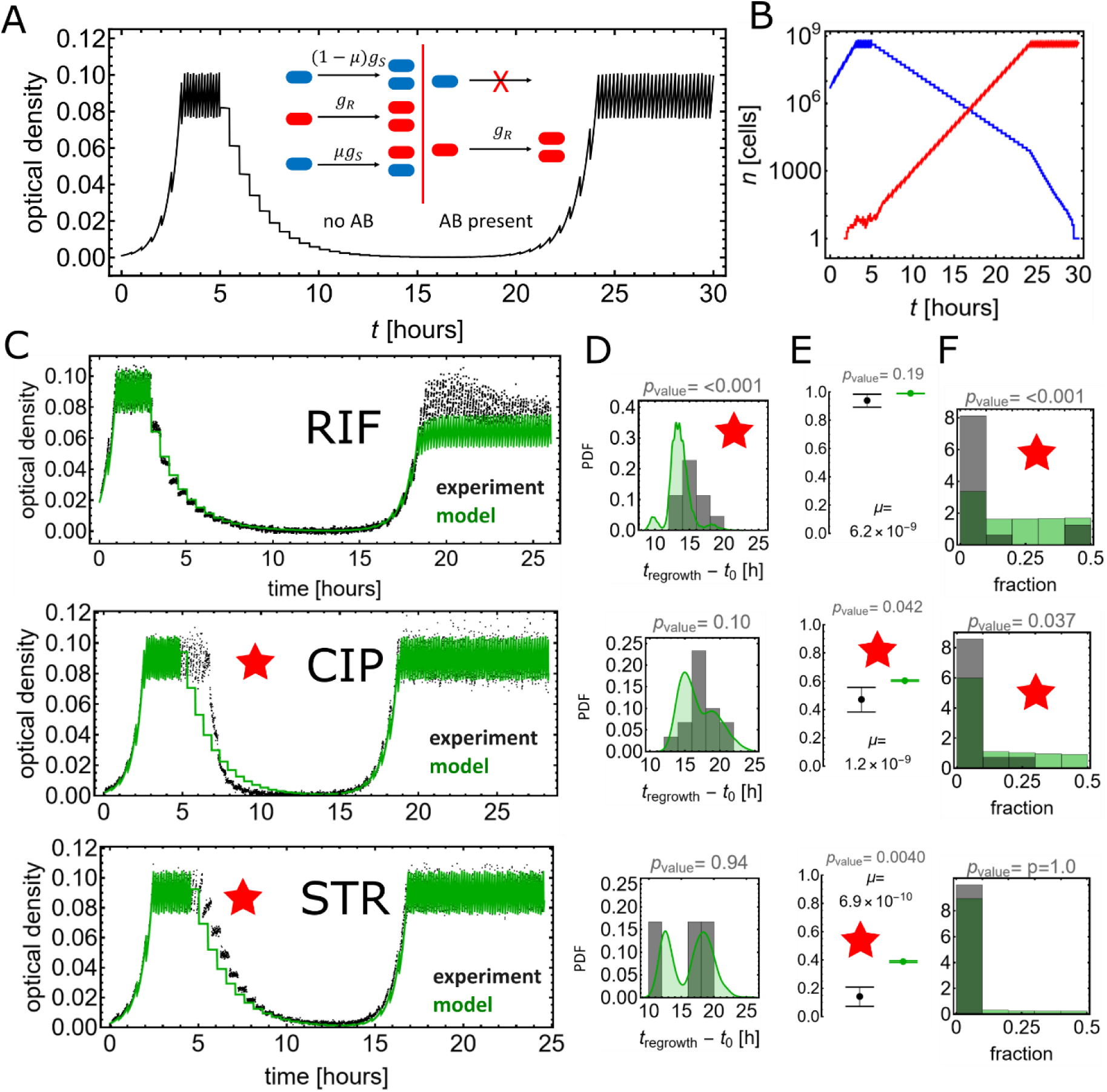
The simple model fails to predict the result of the evolutionary experiment. **(A)** Model definition and an example of optical density (OD) versus time predicted by the model. **(B)** The number of sensitive (blue) and resistant (red) cells versus time in a simulation from panel A. **(C)** Comparison of the model (green) and the experimental (black) OD versus time curves. In each case, only a single experimental and one best-fit theoretical curve has been shown. In the case of RIF we also corrected the experimental data for the contribution of the antibiotic to the optical density, since RIF and its degradation products absorb at OD600 (SI Fig. S1). **(D)** Probability distribution of the regrowth time for cases in which regrowth occurred. **(E)** Probability of regrowth (evolution of resistance). **(F)** Fraction of the least abundant strain at the end of resistant regrowth. Observables that show disagreement between experiment and theory are marked with red stars.

In this model, regrowth upon antibiotic exposure can only occur due to resistant mutations that are already present at the time of antibiotic addition; *de novo* mutations do not occur after antibiotic addition because sensitive bacteria cannot replicate (and hence mutate) in the presence of the antibiotic (Fig. 2B). The number of pre-existing resistant mutants depends on the population size, the number of generations before the antibiotic, and the mutation probability. Since the first two factors are the same for all three antibiotics studied, the probability of regrowth in each case can only be different if *μ* is different for each antibiotic.

To test whether this simple model could quantitatively match our evolutionary experiments, we ran computer simulations using experimentally-derived values of the model’s parameters: the initial number of sensitive cells, the growth rates *g*_S_ and *g*_R_, and the mutation probability *μ*. The initial number of cells was obtained from the initial optical density (Methods), assuming that there were initially no resistant cells. *g*_S_was determined by measuring the growth rate of the sensitive population before the addition of antibiotics and *g*_R_as the growth rate of the population that had re-grown after antibiotic exposure. We also factored in the periodic dilutions in our experimental system. The final parameter required was the mutation probability *μ*; we obtained this by performing mutation fluctuation assays [36] for each of the antibiotics used (Methods, “*Fluctuation test*”, and SI Table S2). Note that, unlike in the bioreactor, bacteria are allowed to enter stationary phase before antibiotic selection in the fluctuation assays.

Figure 2A shows a typical simulated OD versus time curve; this is qualitatively in agreement with the experimental data from Fig. 1C-E. As a first step towards a quantitative comparison between the data and the model, we superimposed simulated OD-vs-time curves on the experimental curves, selecting, for now, only such stochastic realisations of the simulation which matched the experimentally-derived regrowth times. Figure 2C shows an example of an experimental curve and a simulated curve for each antibiotic. Despite the qualitatively good match, some discrepancies are already apparent. In particular, for CIP and STR the sensitive population declines later in response to the antibiotic than predicted by the model.

To further test the model predictions, we estimated the probability that the bacterial population regrows (i.e., evolves resistance), using all our experimental and simulation data, and we also plotted the distribution (across all replicate simulations / experimental runs) of two more quantities: the regrowth time and the fraction of the least abundant strain (FLAS) at the end of resistant regrowth (Fig. 2D-F). The regrowth time is defined as the time since the addition of the antibiotic until the optical density recovers to a value of 0.05 after the sensitive population has stopped growing and been diluted out. FLAS (a value between 0 and 0.5) is obtained by counting the number of colonies of fluorescent/non-fluorescent bacteria after plating an end-point sample on LB agar plates. A monoclonal population would have FLAS=0 whereas a population in which two alleles have equal abundances would have FLAS=0.5.

This quantitative comparison between the simulations and the experimental outcomes further suggests that the simple model, while qualitatively correct, cannot quantitatively predict all aspects of resistance evolution. We consider that the model disagrees with the experiments if the p-value (i.e. the probability that simulation and experimental data are statistically equivalent, for a given observable) is below 0.1. This somewhat conservative threshold p-value ensures that we only consider a model to fit the data if it performs very well. For RIF, the time to regrowth (Fig. 2D) is underestimated by the model and the FLAS is overestimated (Fig. 2F). For CIP, the FLAS and the probability of evolving resistance are overestimated (Fig. 2EF), while for STR, the probability of evolving resistance is overestimated (Fig. 2E).

### Accounting for filamentation of sensitive bacteria upon CIP exposure is necessary to predict the dynamics of CIP resistance evolution

The initial decline of the bacterial population in response to antibiotic is captured correctly by the simple model only for RIF. For RIF, the apparent growth of the population ceases immediately upon exposure, as in the model (Figures 2C and SI Fig. S2). However, for CIP and STR the optical density continues to increase for some time after antibiotic exposure, which is not captured by the model. In the case of CIP, this effect is known to be caused by cell filamentation [37,38], (SI Fig. S4BD): upon exposure cells continue to grow, but do not divide for some time. The continued increase in cell mass is reflected in a continued increase in the optical density even in the absence of cell division.

To model this effect, we added a time-dependent cell density-to-OD conversion factor to the model (see Methods, “*Mathematical modelling*”), which causes the OD to increase for some time post-antibiotic. This improves the model predictions for CIP (SI Fig. S3A and Fig. 4A, column 1). In the case of STR the delayed response to the antibiotic is less significant (SI Fig. S3B) and not caused by filamentation (SI Fig. S4AC). However, we shall still include this factor in all subsequent STR models to ensure this observed (but very small) delay is accounted for in further extensions of the model.

### The model can predict the growth dynamics in the bioreactor in the absence of mutation

After taking into account the initial response of the sensitive population for CIP and STR, quantitative discrepancy still remains between the experimental and predicted data for some of the three measures: regrowth time, regrowth probability, and FLAS. This might suggest that either growth or mutation is incorrectly accounted for by the simple model. To test the correctness of the growth component of the model, we ran “standing variation” experiments in which we inoculated bioreactor cultures with a mixture of resistant bacteria (harvested from previous evolution experiments) and sensitive bacteria in a known proportion (about 1:1000, the exact ratio was determined by plating). A relatively high abundance of the resistant mutant was used to negate any contribution from spontaneous mutations. The model - with the delayed-response for STR and CIP included - provided predictions in very good agreement with the data for all antibiotics (SI Fig. S5). Hence, growth dynamics is correctly represented in the model and the quantitative discrepancies identified earlier likely arise from incorrect modelling of *de novo* mutations and the transition from genetic to phenotypic resistance.

### The generation of CIP and STR resistant clones cannot be predicted using the simple model with a mutation rate determined by a fluctuation test

To test the mutation component of the model, we performed further, “short” experiments to measure the accumulation of *de novo* mutations in the bioreactor. In these experiments, we stopped the bioreactor at the time when the antibiotic would normally have been added in the previous experiments, i.e., two hours after the threshold OD was reached (Fig. 3A).

**Figure 3.**
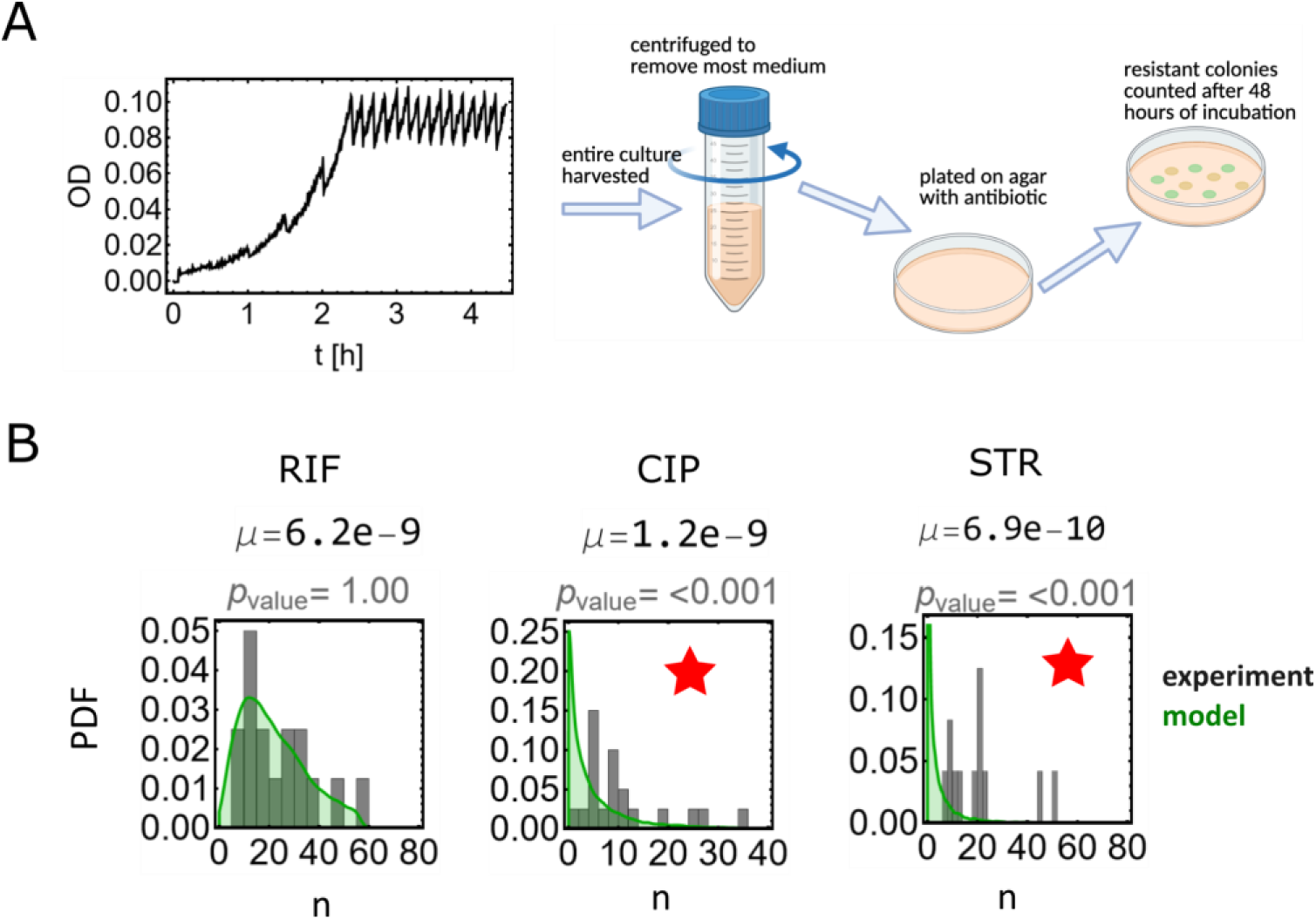
Simple model fails to predict the pre-antibiotic mutant numbers for CIP and STR. **(A)** Schematic of the experiment. Antibiotic-naive bioreactor cultures are spun down and plated on antibiotic-selective agar plates. **(B)** The distribution of the number of resistant colonies obtained experimentally (grey) and predicted by the simple model (green). Disagreement between experiment and theory is marked with a red star.

We then plated all the cells from each culture onto an LB agar plate containing one of RIF, CIP, or STR antibiotics. The number of colonies corresponded to the number of resistant mutant cells that were present in our previous experiments at the moment when the culture was exposed to antibiotics; replicate experiments produced the distribution of mutant numbers. Simulating the same scenario with our model using the mutation rates previously measured in the fluctuation assays, we compared our experimental and simulated mutant number distributions. We found that the simple model could predict the mutant number distribution for RIF, but for CIP or STR the model underestimated the number of resistant bacteria that were present in the bioreactor (Fig. 3B).

This discrepancy suggests that the mutation probability that was measured in the fluctuation assays is not an accurate estimate of the actual mutation probability in the bioreactor. This could be due to the difference in growth conditions: in the bioreactor, cells are maintained in steady exponential growth whereas in the fluctuation assay they experience exponential growth followed by stationary phase. We therefore determined the “true” mutation probability *μ* by fitting the model to the mutant number distributions from the bioreactor experiments (Fig. 3B). For RIF, the estimate for the mutation probability obtained in this way was very similar (*μ* = 8.3 × 10^−9^) to that estimated by the fluctuation assay (*μ* = 6.2 × 10^−9^). However, for CIP and STR the estimated mutation probabilities from the bioreactor data (*μ* = 2.8 × 10^−9^ and 4.7 × 10^−9^ respectively) were significantly higher than the values from the standard fluctuation assays (*μ* = 1.2 × 10^−9^ and 6.9 × 10^−10^, respectively).

Next, we performed simulations of the full experiment, including antibiotic addition, using the “true” mutation rate values listed above. To our surprise, this did not improve the model’s performance. For CIP (Fig. 4A, column 2) the model now overestimated the probability of resistance evolution. The disagreement was even more dramatic for STR, for which the model now also generated incorrect an FLAS prediction (Fig. 4B, column 1).

**Figure 4.**
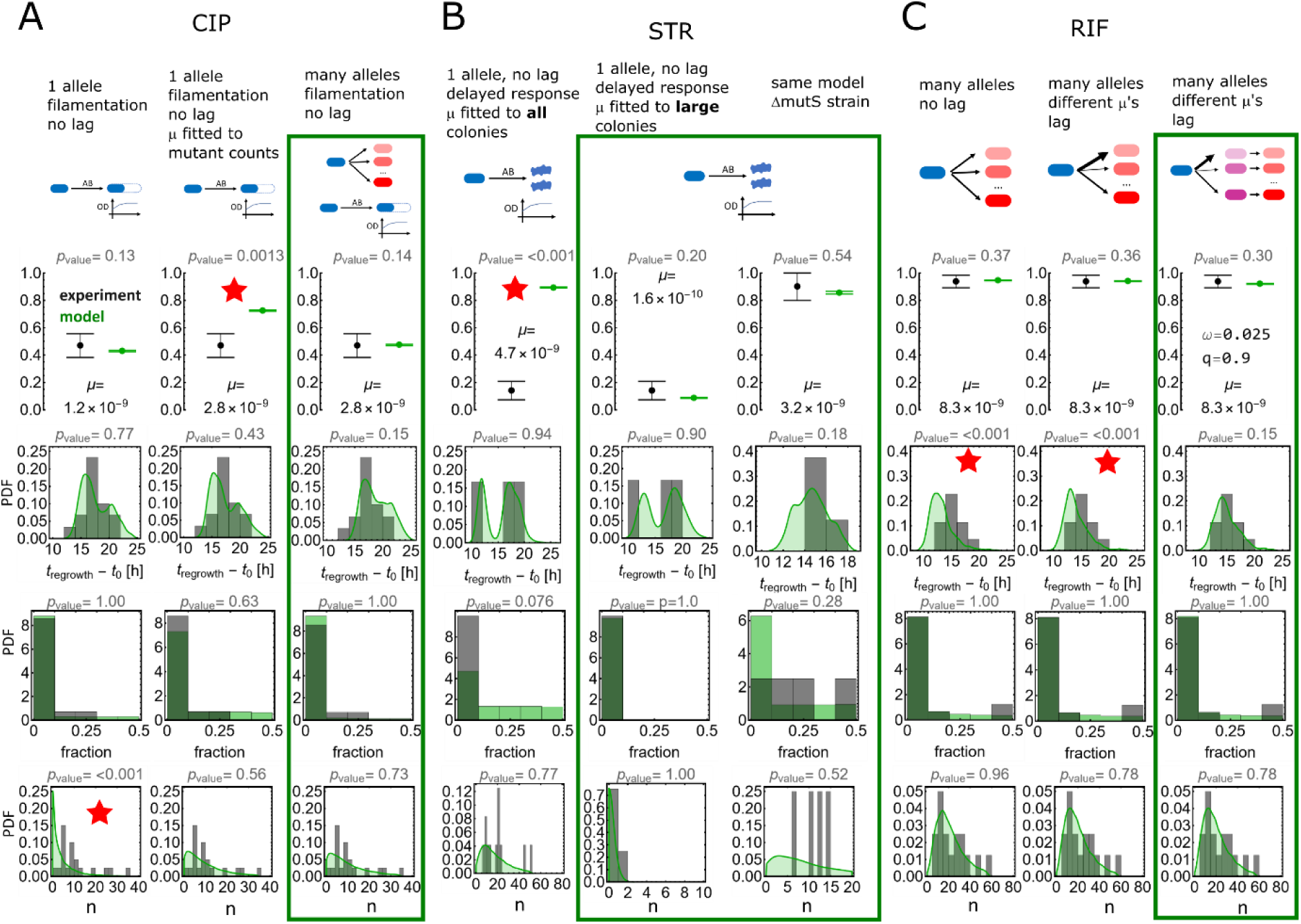
The comparison of different models to the experimental data for ciprofloxacin (A), streptomycin (B) and rifampicin. **(C)**. Rows of plots show: the probability of resistant regrowth, the time to regrowth, the fraction of the least abundant strain at the end of resistant regrowth, and the distribution of the number of resistant mutants prior to antibiotic exposure. Grey and green histograms are the experimental and simulated data, respectively. Model versions for which all experimental and theoretical results statistically agree (p-value >0.1) are surrounded with green rectangles. Observables that show disagreement between experiment and theory are marked with red stars.

### Different phenotypes of resistant alleles need to be taken into account to understand the antibiotic resistance evolution experiments

Our simple model accounts for only one type of resistant mutant (for each antibiotic), but in our STR and RIF experiments we observed differently sized colonies on selective antibiotic-containing agar plates, suggesting the presence of phenotypically different resistant alleles within a single experiment. We therefore decided to include multiple resistant alleles in the model. To do this, we began by characterizing the phenotypes and genotypes of individual clones isolated in our experiments. We included clones both from the long experiments (Fig. 2) and the short experiments (Fig. 3).

Using whole-genome sequencing (WGS), we found that in STR-exposed clones, STR resistance was conferred either by SNPs in *rpsL* (30S ribosomal subunit protein S12, STR target), or by large deletions of 100-150 genes between positions 250 kbp and 400 kbp and no SNPs elsewhere on the MG1655 chromosome. CIP resistance was conferred by SNPs in *gyrA* (DNA gyrase subunit A) or in *gyrB* (DNA gyrase subunit A), both targeted by fluoroquinolone antibiotics. RIF resistance was conferred by SNPs in *rpoB* (beta subunit of bacterial RNA polymerase, RIF target). Supplementary Table 1 lists all these mutations.

With the exception of the large-deletion strains, resistance was always conferred by single mutations in the target of the antibiotic in our experiments. The clones with the large deletion could only be isolated from “short-term” experiments, in which they formed very small colonies on STR agar plates. We concluded that these clones grew too slowly to persist in the long-term evolutionary experiments in the bioreactor. On the other hand, all STR-resistant *rpsL* clones grew at the same rate as the parent strain.

To further investigate the diversity of the resistance mutations to CIP and RIF, we measured the growth rate of all the different clones (identified by WGS) in isolation in the bioreactor in the absence and presence of the antibiotic. The growth rates of RIF and CIP mutants are summarized in Supplementary Table 1.

Since the large-deletion alleles cannot persist in the bioreactor, we concluded that the STR model of bioreactor evolution should take into account only the *rpsL* alleles. Therefore, we obtained a new estimate of the “true” mutation probability for STR by fitting our model to the colony counts from the short STR experiment, taking into account only the normal-sized colonies and ignoring the small colonies. Using this lower mutation rate (*μ* = 1.6 × 10^−10^) the model could now accurately predict the results of evolution in the bioreactor (Fig. 4B, column 2).

Regrowth on STR was very rare in our long experiments, likely due to the rarity of *rpsL* mutations. This limited the size of our dataset for quantitative comparison to the model. To improve the statistics, we decided to repeat all experiments using a strain with a higher mutation rate, which we created by deleting the *mutS* [39] gene. These new experiments showed regrowth in all cultures in long experiments, and short experiments with selection on agar plates yielded both normal and small colony variants, as for the WT strain. Fitting the mutation probability to the colony counts in the short experiments, using only the normal-sized colonies produced a value of *μ* = 4 × 10^−9^. Incorporating this mutation probability into our model including the initially delayed response yielded a single-allele model that could accurately predict all experimental outcomes for the Δ*mutS* strain (Fig. 4B, column 3).

In the case of CIP and RIF, however, our growth rate measurements showed that all alleles could persist in the bioreactor, although the growth rate varied among alleles in both the presence and absence of the antibiotic. Therefore, for CIP and RIF we modified the simple model by including the dynamics of all n alleles (*n*_RIF_ = 21, *n*_CIP_ = 9), using the experimentally measured growth rates. We initially assumed that all alleles are generated with the same mutation probability *μ*/*n*_RIF_or *μ*/*n*_CIP_. We obtained the mutation probability by fitting the model to the mutant number distribution obtained by selection on agar plates in our short experiments, taking all colonies into account. This new model with multiple alleles could now quantitatively predict all experimental outcomes for CIP (Fig. 4A, column 3), suggesting that the inclusion of delayed initial response plus multiple alleles was sufficient to quantitatively explain the dynamics of evolution of resistance to CIP. However, the model still underestimated the time to regrowth for RIF (Fig. 4C, column 1).

Focusing now on RIF, we next questioned our assumption that all alleles are equally likely, since it is known that mutations involving different base pair changes occur with different probabilities in *E.coli* [40,41]. Using the probabilities obtained from Refs. [40,41] in the multiple allele model improved its performance for the distribution of RIF regrowth times, but the p-value for the comparison between model and experiment was still below 0.001, suggesting significant disagreement (Fig. 4C, column 2).

### The addition of phenotypic delay can explain the dynamics of RIF resistance evolution

Thus far our models assumed that a mutation immediately generates a fully resistant cell that grows with the population growth rate measured for that allele. However, it is known that some antibiotics (including the ones we use here) exhibit a phenotypic delay in the emergence of resistance [42,43]. This occurs in cases where antibiotic sensitivity is phenotypically dominant over resistance [44,45]. Immediately after the genetic mutation, a cell contains both resistant and sensitive variants of the target protein; the sensitive variants must be diluted out by growth and cell division before the cell assumes a fully resistant phenotype. Thus, there is a time delay before newly generated mutants become fully resistant. Although our experimental setup does not allow for direct measurement of this “phenotypic delay”, we hypothesized that it might account for the remaining difference between the model predictions and our RIF experimental data.

To explore this, we added a simple phenotypic delay to our model in the form of an intermediate, semi-resistant phenotype. This phenotype is created from the sensitive strain with mutation probability *μ* but in the presence of the antibiotic its growth rate is only a fraction *q* of the growth rate of fully resistant phenotype. The intermediate phenotype switches to the fully resistant phenotype with rate *ω* (≫ *μg*_S_), with 1/*ω* representing the average length of phenotypic delay. This model is a crude approximation of more accurate mechanistic models of phenotypic lag [42,43].

To find the best values for *ω* and *q*, we fitted the model to all our experimental observables from the long, evolutionary bioreactor runs (Methods, “*Optimal phenotypic lag parameters*”). Adding a long but weak phenotypic delay (1/*ω* ∼ 40 h, *q* = 90% of the resistant growth rate) to the multiple allele model of RIF evolution improved its fit to the experimental data (Fig. 4C, column 3), such that the model could now reproduce the distribution of regrowth times with a p-value of above 0.05 (both with and without the inclusion of different mutation rates for different alleles, cf. SI Fig. S6). Although the CIP data could be explained (with p-values above 0.05 for all experimental observables) without a phenotypic delay, models with a short but phenotypically strong delay (1/*ω* = 2 … 3 h, *g* = 0. . .30%) also improved the agreement for this antibiotic (SI Fig. S7). We stress that the numerical values of the delay are difficult to interpret biologically because our model is not mechanistically correct; rather, it should be considered a simple, effective model of the emergence of phenotypic resistance from genetic mutations.

## Discussion

Simple birth-death-mutation models are widely used to predict the dynamics of antibiotic resistance evolution, but it is not clear how quantitatively accurate their predictions are. In this work we performed exhaustive tests of our ability to model resistance evolution, using a quantitative dataset from a bioreactor setup that was optimized to facilitate the comparison with models, in that bacteria are kept in steady state growth and cell density and growth rate can be accurately measured, as well as the timing of resistance emergence. Our results show that although a simple birth-death-mutation model can accurately predict some aspects of the experimental evolution of antibiotic resistance, it fails to quantitatively predict all experimental outcomes. We find that the model fails for reasons that are different depending on the antibiotic used. This in turn provides insight into the mechanism of AMR evolution for each antibiotic.

The model with a single resistant allele, a mutation probability obtained from a standard mutation fluctuation test and accounting for a delayed initial response due to filamentation of the sensitive population can actually predict the outcome of resistance evolution to CIP: the probability of regrowth, and the time to regrowth as well as the fraction of the least-abundant allele. However, this model fails to predict the number of resistant mutants present before antibiotic exposure. Explicitly including different resistant alleles with experimentally measured growth rates is necessary to fully explain all the data, including the mutant number distribution.

In the case of STR, for our wild-type *E. coli* strain the probability of resistance is very low (Fig. 2D), and thus we had fewer experimental datasets to compare the model to. However, we find that neither the mutation rate obtained from a standard mutation fluctuation test, nor a revised mutation rate obtained from counting colonies isolated on STR plates from growth in the bioreactor, accurately predicts the regrowth probability. This is because mutations generate an abundance of slowly-growing resistant variants conferred by large genomic deletions but relatively few fast-growing alleles in the STR target *rpsL*. Interestingly, large deletions detected by WGS always included the *hemB* gene (delta-aminolevulinic acid dehydratase), and *ΔhemB* strains are known to produce the “small-colony variant” we observed. Although the *ΔhemB* variants are known to persist in infections [46], their slow growth causes them to be washed out of the bioreactor and thus do not contribute to the evolution of resistance. In contrast, in the standard fluctuation assay, bacteria are grown in batch cultures and not being diluted out, and thus *rpsL* mutations which are STR-dependant can be selected for [33,47].

Finally, for RIF, evolution of resistance occurred in almost all long-term experiments, thanks to the large target size (many different SNPs in *rpoB* can confer resistance, see Table S1) [34]. However, including the 21 different alleles of *rpoB* that we identified in our experiments together with their different mutation probabilities was not enough to fully explain all the experimental outcomes. Of course, our model with multiple alleles does not take into account all the possible *rpoB* mutations that can lead to RIF resistance [48,49], but considering the diversity in growth rates already included it is unlikely that including more alleles would improve the model. In particular, the model underestimated the time it takes for a resistant population to regrow in the bioreactor (Fig. 4C). We therefore explored the phenomenon of phenotypic delay [43]. It has been experimentally demonstrated that some resistance mutations in *rpoB* as well as *rpsL* and *gyrA* cause phenotypic delay due to polyploidy, in which the multiple chromosomes in fast growing cells have to be diluted out before phenotypic resistance to the antibiotic can occur [42]. We modelled this in a very simplistic way by introducing an intermediate phenotype occurring upon mutation, which grows in the presence of the antibiotic at a slower rate than the resistant phenotype, and which slowly transitions to full resistance. In reality the delay will depend on cells’ history: the number of resistant/sensitive copies of the target gene, the number of resistant/sensitive target molecules, position in the cell cycle, and other factors. Nevertheless, this simple model substantially improved the agreement with the RIF data.

Our findings for RIF do not constitute a proof of phenotypic delay, nor do they explain its mechanism. Nevertheless, the model with phenotypic delay can be treated as an effective model of resistance to RIF that explains all our experimental results. Our results do not rule out phenotypic delay in the case of CIP and STR; they only show the delay is not required to model our experimental data. This could be due to the delay being very short, or not contributing to the observables we chose to look at. For example, a delay that would cause bacteria containing sensitive target molecules not to grow at all (*q* = 0) would most likely manifest itself as a lower effective mutation probability in both the fluctuation test and the short-term bioreactor experiment. Detecting this type of delay would require a different experimental approach to that used here, possibly involving single-cell imaging [42].

Interestingly, the mutation rates for STR and CIP that we measured in the standard fluctuation assays were significantly lower than the mutation rates we obtained from the bioreactor experiments. This suggests that continuous steady state growth may contribute to the increase in the emergence of phenotypically resistant alleles. One possible explanation is that in the bioreactor, cell lineages may have more time (generations) to overcome a phenotypic delay that would otherwise prevent growth upon selection. That is, in a batch culture in the standard fluctuation assay, most mutations would likely occur at the highest cell densities when nutrients are almost depleted, allowing for few generations after the mutation to dilute out sensitive chromosomes and proteins.

Taken together, our results show that a simple birth-death-mutation model cannot predict all the features of the dynamics of resistance evolution in a bioreactor, even though our experiments were designed to be as simple as possible (i.e., with as few unmeasurable/ uncontrolled variables as possible). It should be noted that whether the simple model can quantitatively predict experimental outcomes depends on which outcome is looked at, and which outcomes can be accurately predicted, and the necessary modifications to the model, depend on which antibiotic is used. Thus, we conclude that, when considering the outcome of treating a bacterial population with an antibiotic, it is important to take into account the initial response of the population, the fitness of different resistant alleles, and phenotypic delay. Not accounting for these factors can lead to models that appear correct at a qualitative level, but fail to quantitatively predict bacterial growth and dynamics, which compromises their usefulness for modelling personalized antibiotic therapy [50].

## Materials and Methods

### Bioreactor setup

We use a bioreactor system described before in Ref. [31] that maintains approximately constant turbidity of the culture by repeatedly diluting it with fresh medium using a negative feedback loop. This keeps the bacteria in the exponential growth phase in the absence of antibiotics. Our system is close to a turbidostat, however the turbidity is allowed to fluctuate in a range between OD=0.075 and OD=0.1 to enable fitting an exponential function to segments of OD vs time curve; this gives an accurate estimate of the growth rate.

The system consists of four replicate cultures with an approximate working volume of 25 mL each. Each bottle has two inlets: one for the LB medium, the other for the LB+antibiotic, an outlet for removing excess liquid, a sampling port, a tube delivering air for culture aeration, a port for a spectrophotometer, and a resistance-based liquid level sensor. A distinguishing feature of our setup are custom-made syringe pumps and valves which enable us to precisely (<1%) control the volume of injected and pumped-out media. Each bottle is equipped with a miniature spectrophotometer to measure the optical density. This is achieved by aspirating each culture every 10 s into a specially designed glass cuvette at the top of the bottle, passing light from a narrow-beam LED (wavelength = 600 nm) through it, and measuring the intensity of the incoming and transmitted light.

All cultures are connected to an air pump for aeration, and stirred vigorously using magnetic beads at the bottom of each bottle. An additional sample tube with a syringe at the end is used for inoculation and sampling. The culture bottles as well as the pumps and the electronics are placed in a temperature-controlled incubator with fans that ensure uniform temperature of all cultures.

The bioreactor is controlled by a PC using custom-made software that monitors the optical density, temperature, and controls the injection of LB and antibiotics. Dilution occurs whenever the OD reaches OD=0.1 or after 30mins, whichever happens first. Each dilution replaces 25% of the culture volume with fresh medium. This guarantees that only cells with growth rate *g* > (−log 0.75)/ 0.5 = 0.58 h^-1^ (doubling time < 72 min) can persist in the culture. The software automatically calculates how much antibiotic to add to achieve a pre-determined, time-dependent concentration in the culture. The actual concentration of the antibiotic is not monitored.

Prior to the experiment, culture bottles are removed from the system and autoclaved, and (after re-attaching the bottles) the system is flushed with bleaching agent (PRESEPT, 5 g / 1 L water) and three times with media to dilute out the bleach. This approach effectively sterilizes the equipment (no growth over many days in test runs with no inoculum). The incubator is switched on and, after the system has reached 37°C, each culture is inoculated with 1ml of bacterial suspension.

### Evolution experiments

Overnight cultures of *E. coli* AD30 and EEL02 were diluted 1:200 in 10mL each of fresh medium and pre-grown shaking at 37°C for 1 h before being mixed 1:1 and 1mL of this mixture was injected into each bottle. The inoculum was also diluted 1: 10^4^ into PBS and plated or spotted onto LB agar (100 μL/plate) and inoculated overnight at 37°C for CFU counts. The software was set to dilute each culture when OD600 of 0.1 was reached, or every 30 min. In the long-term experiments the antibiotic was added 2 hours after the optical density reached OD=0.1 for the first time, and then maintained at the same concentration throughout the rest of the experiment. At the end of long-term experiments, all cultures where resistant regrowth had occurred were sampled by diluting the culture 1:10^4^in PBS and spotted or plated on LB agar for CFU counts (100 μL/plate). A sample of each resistant population was also frozen in –80C in 15% glycerol. In the short-term experiments in which pre-antibiotic resistant mutants were counted on antibiotic-agar plates, the experiments were stopped approximately 2 h after the optical density reached OD=0.1 for the first time. The times were not exactly 2 h because all 4 cultures had to be stopped at the same time but not all cultures reached OD=0.1 at exactly the same time. The entire cultures were poured into Falcon tubes kept on ice to prevent further cell division before plating. A small sample (10 μL) from each culture was diluted 1: 10^4^into PBS before being spotted onto LB agar for CFU count and incubated at 37°C overnight, and the rest of the culture were centrifuged at 4C, 4700 RPM for 45 min. Most of the supernatant was carefully poured off and the pellet was resuspended in the remaining medium. The resuspended pellet was plated on LB agar plates containing the antibiotic used for selection. The plates were then incubated at 37°C for 48 hours to allow for growth of all resistant mutants present in the culture.

“Standing variation” experiments used a 1:1000 mixture (v/v) of overnight cultures of resistant:sensitive bacteria, pre-grown outside the bioreactor to make sure that the resistant culture as well as the sensitive wild type were growing exponentially upon inoculation, as confirmed by measuring OD over time. To get the actual initial ratio of resistant to sensitive cells, the inoculums were diluted 10x and spotted onto LB agar with the antibiotic for resistant cell count, and diluted 1: 10^4^ and spotted onto LB agar for total cell count, and then incubated overnight at 37°C.

To measure the growth rates of the resistant mutants in the bioreactor in the absence and presence of the antibiotic, each identified allele was grown overnight, diluted 1:200 in fresh LB and pre-grown for 1 h shaking at 37°C. 1 mL was then inoculated into the bioreactor. The antibiotic relevant for the resistance was added 2 h after the optical density reached OD=0.1. In the cases where the alleles grew too slowly in the bioreactor, the time between dilutions was changed to 2 h instead of 30 min to enable the slow-growing alleles to persist in the culture.

### Fluctuation tests

A single bacterial colony was picked and inoculated in 1 mL of LB medium and grown overnight to stationary phase. This stationary culture was diluted 10^6^ times and cultured again for 24 hours to reach the stationary phase. This culture was diluted again 10^6^ times, divided into many small cultures. A small volume of the diluted culture was also plated onto non-selective agar to estimate the initial number of colony forming units (CFUs). All the replicate small cultures were again incubated at 37°C for 24 hours, to ensure they reached stationary phase. Then a few cultures were diluted 10^6^ times and plated on LB agar plates to estimate the final density of cells. The remaining cultures were plated onto selective agar plates and incubated for 48 h, and the number of CFUs was counted. All results can be found in Supplementary Table 2.

### Bacterial strains

We used a derivative of the *E.coli* K12 strain MG1655 with a *fimA* deletion (AD30) [31], and an isogenic strain with constitutive YFP expression and chloramphenicol resistance (EEL02). The *fimA* deletion prevents biofilm formation in the bioreactor (as opposed to MG1655), probably due to the lack of functional fimbriae. The cassette with constitutive YFP expression and chloramphenicol resistance were moved from strain RJA002 [51] using P1 transduction [52]. In the STR experiment with an increased mutator rate we used two isogenic mutator strains where *mutS* was deleted using plasmid mediated gene replacement [53], using the following primers to amplify the upstream and downstream regions of the *mutS* gene: mutSup_fwd; AAAAACTGCAGAAACAACGCCTCGTAATGCT, mutSup_rev; CGGGAATTGTTAGGGGTTATGTCCGGT, mutSdown_fwd: GGACATAACCCCTAACAATTCCCGATA and mutSdown_rev: AAAAAGTCGACCACCGCTGGCAAGCCACATA, and using crossover PCR to anneal the regions before ligation into the pTOF24 plasmid used for the gene deletion. We deleted *mutS* in AD30 and an isogenic strain (EEL03) with constitutive CFP expression and ampicillin resistance, the cassette with CFP and ampicillin resistance were moved from strain RJA003 [51], using P1 transduction [52].

### Growth conditions

All experiments were performed in LB broth (Miller), at 37°C, and on LB (miller) agar plates with 1.5% agar-agar. Antibiotic stocks were prepared as follows; 50 mg/ml rifampicin in DMSO, 10 mg/ml ciprofloxacin hydrochloride in water, and 100 mg/ml streptomycin in water. We used concentrations of 100 μg/ml of rifampicin and streptomycin and 100 ng/ml ciprofloxacin in all experiments.

### Whole genome sequencing to identify resistance conferring mutations

Chromosomal DNA was extracted from each clone using the Monarch Genomic DNA Purification Kit (New England Biolabs). DNA libraries were prepared using the Rapid Sequencing DNA Kit with 96 barcodes (SQK-RBK110.96 or SQK-RBK114.96). The libraries were sequenced using the MinION Mk1B device (Oxford Nanopore), with R9.4.1 or R10.4.01 flow cells. The sequence data for all sequenced alleles has been deposited in the European Nucleotide Archive at EMBL-EBI under accession number PRJEB86481.

Basecalling, barcode splitting and aligning the resulting FASTQ files to the reference genome (*Escherichia coli* str. K-12 substr. MG1655, reference genome assembly GCF_000005845.2_ASM584v2) was done using guppy (Oxford Nanopore Technologies plc., version 6.5.7+ca6d6af). Variant calling was performed with freebayes (v1.3.6) [54] using the default parameters, except that the minimum coverage was set to 5. To detect large deletions, we used delly (v1.1.5) [55] as well as visual inspection of genome coverage using Geneious (https://www.geneious.com/). Since Nanopore sequencing is known to generate many artefacts, especially in regions rich in k-mers [56], we treated all detected variants containing k-mers with *k* > 3 as artefacts. To further reduce the probability of artefacts when detecting SNPs, we focused only on genes with known relevance for RIF, CIP, and STR resistance: *gyrA*, *gyrB*, *parC*, *marR*, *marA*, *marB*, *rpsL*, *rpoB*. Occasional SNPs detected in other regions showed no consistency between the altered region and the antibiotic used, and did not repeat in samples treated in the same way. We therefore considered such SNPs to represent sequencing artefacts.

### Bioreactor data analysis

All analysis has been done using Wolfram Mathematica notebooks.

#### Growth rate

For each OD-vs-time curve we obtained a sequence of growth rates {*g*_*i*_} corresponding to time periods between consecutive dilutions by fitting a piece-wise exponential function with a background correction

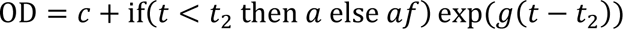

to every two consecutive dilution cycles *t*_1_ → *t*_2_ and *t*_2_ → *t*_3_. This allowed us to a more unambiguous estimation of the background correction *c* than it would be possible by fitting a single exponential function to a single cycle. The approach was tested using synthetic data. The accurate determination of the dilution factor *f* = 0.75 plays an important role here, hence we performed experiments with the methylene blue dye subject to periodic dilutions to make sure the dilution factor was 0.75 in each culture as assumed above.

#### Growth rate of the sensitive population

We took all {*g*_*i*_} that came from dilution cycles before the antibiotic and in which the minimum OD was at least 0.05.

#### Growth rate of the resistant population

We took all {*g*_*i*_} from the regrowth phase which had the minimum OD of at least 0.05.

#### Initial optical density at t = 0

Since the culture was initially very diluted and measuring its OD would not be accurate, we obtained OD_initial_by extrapolating backwards from the growth phase with OD>0.075 using the growth rate determined as described above, and assuming exponential growth since the beginning of the experiment. This gave OD_initial_ values between 0.002 and 0.005, in agreement with what was expected based on the volume and density of the inoculum as well as the direct measurement of OD_initial_at *t* = 0.

#### Conversion factor *Q* between the optical density and bacterial density

To establish the proportionality coefficient between bacterial and optical density (no. of cells/ml per OD=1), we collected samples from a single bioreactor run (4 cultures) corresponding to the steady-state exponential growth phase (about 5000 s since the OD reaching OD=0.1 for the first time). The samples were diluted 1: 10^4^ times and plated (100 ml) on LB agar plates (two replicates for each culture). To maximize plating efficiency, we spot-plated each 100 ml, i.e., we touched the agar with a pipette tip at tens of positions, each time expressing only a small volume of the 100 ml total volume. We verified by comparing colony counts with counting bacteria from the same cultures under the microscope that this technique was much more reliable than expressing the entire volume at once and spreading it using a sterile spreader. The corresponding optical densities and cell counts were (OD, CFUs) = [[0.0823, 196 + 176], [0.0929, 155 + 199], [0.0866, 216 + 194], [0.0939, 206 + 183]], from which we calculated the conversion factor to be 2.15 × 10^8^ bacteria/unit OD. In the simulations, we used a rounded-down number *Q*^−1^ = 2 × 10^8^ bacteria/OD.

#### Mathematical modelling

We used the generalised version of the birth process [23] with two species: sensitive and resistant cells, and with mutations. Sensitive cells divide with rate *g*_S_in the absence of antibiotics, and do not replicate when the antibiotic is present. Replication results in a new sensitive cell with probability 1 − *μ*, or with a resistant cell with probability *μ*. Resistant cells divide with rate *g*_R_(same in the presence/absence of the antibiotic).

To simulate bioreactor experiments, the number of sensitive bacteria *n*_S_ is set to the number calculated from the initial optical density as explained above, and the number of resistant bacteria *n*_R_ is set to zero. Growth is simulated using an approximate stochastic simulation protocol - the “tau leaping algorithm” [57] with a fixed time step of 1/64 h.

Dilution is implemented by generating new *n*_S_′, *r*_R_′ from binomial distributions with means *n*_S_, *r*_R_ and the success rate *f* = 0.75 equals the fraction of the volume retained after a single dilution step.

To model filamentation (CIP) or a more general delayed response to the antibiotic (STR), we used a time-varying conversion factor between the number of cells and the optical density: OD = OD_R_ + OD_S_

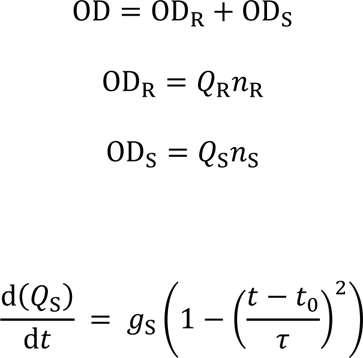

where *Q*_*S*_(0) = *Q*_*R*_ = 1/(2 × 10^7^), and *τ* is set to 2 h for CIP and 0.66 h for STR to account for the observed time delay (SI Fig. S2).

To model different mutation probabilities, we first estimated the relative probability *P*_allele *i*_ of occurrence of different SNPs found in our evolutionary experiments. We then modified the model so that, while the total mutation probability to resistance was still *μ*, a specific allele *i* would be selected with probability *μP*_allele *i*_.

To account for RIF contribution to the OD, we modelled RIF and its decay product rifampicin quinone (RIFO) using the following differential equations

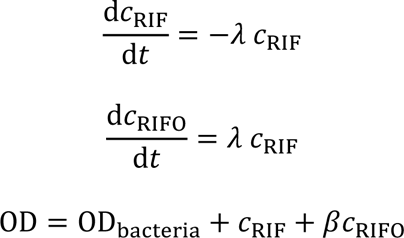

Dilution was modelled as follows: *c*_RIF_ → *f c*_RIF_ + (1 − *f*)*c*_0,RIF_, *c*_RIFO_ → *fc*_RIFO_ The model has three parameters: *c*_0,RIF_which describes the theoretical OD contribution if the entire culture was replaced by RIF from the reservoir, *β* which describes by how much more the decay product contributes to the OD compared to RIF, and *λ* which is the RIF decay rate. We first estimated *β* by fitting this model to an experiment in which we added RIF to the culture in the same way as in the long-term evolutionary experiment but without any bacteria present. The *λ* rate could in principle depend on the chemical composition of the culture, in particular its pH, and temperature. Similarly, *c*_0,RIF_could differ for different experiments since RIF may have decayed already in the reservoir to an unknown extent. This is not a problem from the RIF antimicrobial activity standpoint because the RIF quinone is also active against bacteria, but it means each replicate RIF experiment may have a slightly different background OD after the first RIF injection. Therefore, to obtain *λ* and *c*_0,RIF_ we fitted the RIF decay model to experimental long-term OD vs time curves in the regime in which the curve was close to zero (almost no bacteria in the culture), and the only contribution to OD was due to RIF and RIFO. This gave the value of *λ* between 0.15 and 0.2 h^-1^ for different experiments, and *c*_0,RIF_ between 0.016 and 0.032. These values were then used to subtract the RIF contribution each individual OD-vs-time curve.

All models have been implemented in C++.

### Statistical tests

To compare experimental data and model predictions we used Kolmogorov-Smirnov test (*KolmogorovSmirnovTest* in Wolfram Mathematica) for quantities such as the regrowth time, the fraction of the least-abundant strain, and the number of resistant mutants. We used Fisher exact test (our own implementation in Mathematica) for the regrowth probability. All these tests return a p-value which is what we use to determine whether a model is a good fit to the data (p-value above 0.1) or not.

### The mutation probability from fluctuation test data

We used the Lea and Coulson model (birth process with mutations) [23] to determine *μ* from the fluctuation test data. Specifically, we used *P*(*m*; *μ*), the probability of finding *m* mutants assuming the mutation probability *μ* per replication, to calculate the posterior probability distribution for a vector of *N* measurements {*m*_1_, . . ., *m*_*N*_}:

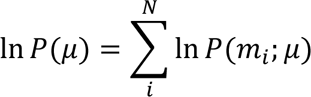

and find a value of *μ* which maximized this probability. To calculate *P*(*m*; *μ*) we used the recursive relation [36].

### Optimal mutation probability *μ* from the mutant distribution data

We searched for a value of *μ* for which the p-value of the Kolmogorov-Smirnov test comparing the experimental and simulated distributions of the number of mutants before the antibiotic was maximized. We wrote a Wolfram Mathematica function which returned this p-value using a simulated distribution from 300 independent simulations, for a given value of *μ*, and used this in a minimization routine (“golden-section search”), suitably bracketing the optimum *μ* to a biologically-realistic range.

### Optimal phenotypic delay parameters

We proceeded similarly to when finding the optimal *μ*, but this time we maximized the expression:

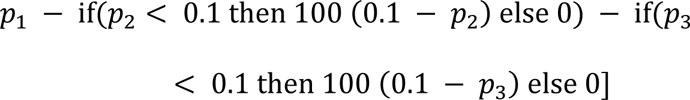

where *p*_1_, *p*_2_, *p*_3_ are p-values of KS tests comparing the simulated and experimental observables: the regrowth probability, the distribution of regrowth times, and the distribution of the least-frequent allele. The function significantly penalized any p-value larger than 0.1, thus forcing such parameter values that would lead to all p-values below 0.1 if at all possible. Since the optimization problem was now two-dimensional and the function could in principle have many local minima, we used a brute-force approach: we calculated the fitting function for a range of *μ* and *ω* values regularly spaced on a grid, with a suitably small step in each direction, and biologically realistic boundaries. See the Mathematica code for more details.

### Data and code

All code (C++ code for simulating the model, Mathematica notebooks for data analysis and plotting) and bioreactor data are available on GitHub: https://github.com/Dioscuri-Centre/AMR_bioreactor. DNA sequences are available from the European Nucleotide Archive at EMBL-EBI under accession number PRJEB86481.

## Acknowledgments

This work was funded through Dioscuri, a programme initiated by the Max Planck Society, jointly managed with the National Science Centre in Poland, and mutually funded by Polish Ministry of Science and Higher Education and German Federal Ministry of Education and Research, grant no. UMO-2019/02/H/NZ6/00003. RJA was supported by the European Research Council under Consolidator grant 682237 EVOSTRUC and by the Deutsche Forschungsgemeinschaft (DFG) under the Excellence Cluster Balance of the Microverse (EXC 2051 – Project ID 390713860). We thank Ilyas Djafer-Cherif for his help with analysing WGS data. We also thank Helen Alexander, Nikola Ojkic, Martin Carballo-Pacheco, Michael Nicholson, and the whole Dioscuri Centre team for many helpful discussions and feedback on various aspects of this project. Figure 3A was created with BioRender.com

## References

1. Robinson TP, Bu DP, Carrique-Mas J, Fèvre EM, Gilbert M, Grace D, et al. Antibiotic resistance is the quintessential One Health issue. Trans R Soc Trop Med Hyg. 2016;110: 377–380. doi:10.1093/trstmh/trw048

2. Aslam B, Wang W, Arshad MI, Khurshid M, Muzammil S, Rasool MH, et al. Antibiotic resistance: a rundown of a global crisis. Infect Drug Resist. 2018;Volume 11: 1645– 1658. doi:10.2147/IDR.S173867

3. Murray CJL, Ikuta KS, Sharara F, Swetschinski L, Robles Aguilar G, Gray A, et al. Global burden of bacterial antimicrobial resistance in 2019: a systematic analysis. The Lancet. 2022;399: 629–655. doi:10.1016/S0140-6736(21)02724-0

4. Wong A, Rodrigue N, Kassen R. Genomics of Adaptation during Experimental Evolution of the Opportunistic Pathogen Pseudomonas aeruginosa. Guttman DS, editor. PLoS Genet. 2012;8: e1002928. doi:10.1371/journal.pgen.1002928

5. Zhang Q, Lambert G, Liao D, Kim H, Robin K, Tung C, et al. Acceleration of Emergence of Bacterial Antibiotic Resistance in Connected Microenvironments. Science. 2011;333: 1764–1767. doi:10.1126/science.1208747

6. Jansen G, Barbosa C, Schulenburg H. Experimental evolution as an efficient tool to dissect adaptive paths to antibiotic resistance. Drug Resist Updat. 2013;16: 96–107. doi:10.1016/j.drup.2014.02.002

7. Jahn LJ, Munck C, Ellabaan MMH, Sommer MOA. Adaptive Laboratory Evolution of Antibiotic Resistance Using Different Selection Regimes Lead to Similar Phenotypes and Genotypes. Front Microbiol. 2017;8: 816. doi:10.3389/fmicb.2017.00816

8. Toprak E, Veres A, Yildiz S, Pedraza JM, Chait R, Paulsson J, et al. Building a morbidostat: an automated continuous-culture device for studying bacterial drug resistance under dynamically sustained drug inhibition. Nat Protoc. 2013;8: 555–567. doi:10.1038/nprot.2013.021

9. Spagnolo F, Rinaldi C, Sajorda DR, Dykhuizen DE. Evolution of Resistance to Continuously Increasing Streptomycin Concentrations in Populations of Escherichia coli. Antimicrob Agents Chemother. 2016;60: 1336–1342. doi:10.1128/AAC.01359-15

10. Pinheiro F, Warsi O, Andersson DI, Lässig M. Predicting trajectories and mechanisms of antibiotic resistance evolution. arXiv; 2020. doi:10.1101/2020.07.02.184622

11. Austin D, Anderson R. Studies of antibiotic resistance within the patient, hospitals and the community using simple mathematical models. Philos Trans R Soc Lond B Biol Sci. 1999;354: 721–738.

12. Wu Y, Saddler CA, Valckenborgh F, Tanaka MM. Dynamics of evolutionary rescue in changing environments and the emergence of antibiotic resistance. J Theor Biol. 2014;340: 222–231. doi:10.1016/j.jtbi.2013.09.026

13. Nissen-Meyer S. Analysis of Effects of Antibiotics on Bacteria by Means of Stochastic Models. Biometrics. 1966;22: 761. doi:10.2307/2528073

14. Torella JP, Chait R, Kishony R. Optimal drug synergy in antimicrobial treatments. PLoS Comput Biol. 2010;6: e1000796.

15. Bonhoeffer S, Lipsitch M, Levin BR. Evaluating treatment protocols to prevent antibiotic resistance. Proc Natl Acad Sci. 1997;94: 12106.

16. Nichol D, Jeavons P, Fletcher AG, Bonomo RA, Maini PK, Paul JL, et al. Steering evolution with sequential therapy to prevent the emergence of bacterial antibiotic resistance. PLoS Comput Biol. 2015;11: e1004493.

17. Brauner A, Balaban NQ. Quantitative biology of survival under antibiotic treatments. Curr Opin Microbiol. 2021;64: 139–145. doi:10.1016/j.mib.2021.10.007

18. Sun L, Ashcroft P, Ackermann M, Bonhoeffer S. Stochastic Gene Expression Influences the Selection of Antibiotic Resistance Mutations. Mol Biol Evol. 2020;37: 58–70. doi:10.1093/molbev/msz199

19. Alexander HK, MacLean RC. Stochastic bacterial population dynamics restrict the establishment of antibiotic resistance from single cells. Proc Natl Acad Sci. 2020;117: 19455–19464. doi:10.1073/pnas.1919672117

20. Coates J, Park BR, Le D, Şimşek E, Chaudhry W, Kim M. Antibiotic-induced population fluctuations and stochastic clearance of bacteria. eLife. 2018;7: e32976. doi:10.7554/eLife.32976

21. Pena-Miller R, Laehnemann D, Jansen G, Fuentes-Hernandez A, Rosenstiel P, Schulenburg H, et al. When the Most Potent Combination of Antibiotics Selects for the Greatest Bacterial Load: The Smile-Frown Transition. PLOS Biol. 2013;11: e1001540. doi:10.1371/journal.pbio.1001540

22. Muetter M, Angst DC, Regoes RR, Bonhoeffer S. The impact of treatment strategies on the epidemiological dynamics of plasmid-conferred antibiotic resistance. Proc Natl Acad Sci. 2024;121: e2406818121. doi:10.1073/pnas.2406818121

23. Lea DE, Coulson CA. The distribution of the numbers of mutants in bacterial populations. J Genet. 1949;49: 264. doi:10.1007/BF02986080

24. Kendall DG. Birth-and-Death Processes, and the Theory of Carcinogenesis. Biometrika. 1960;47: 13–21. doi:10.2307/2332953

25. Nicholson MD, Antal T. Universal Asymptotic Clone Size Distribution for General Population Growth. Bull Math Biol. 2016;78: 2243–2276. doi:10.1007/s11538-016-0221-x

26. Gralka M, Fusco D, Martis S, Hallatschek O. Convection shapes the trade-off between antibiotic efficacy and the selection for resistance in spatial gradients. Phys Biol. 2017;14: 045011. doi:10.1088/1478-3975/aa7bb3

27. Greulich P, Waclaw B, Allen RJ. Mutational pathway determines whether drug gradients accelerate evolution of drug-resistant cells. Phys Rev Lett. 2012;109: 088101. doi:10.1103/PhysRevLett.109.088101

28. Nande A, Hill AL. The risk of drug resistance during long-acting antimicrobial therapy. Proc R Soc B Biol Sci. 289: 20221444. doi:10.1098/rspb.2022.1444

29. Igler C, Rolff J, Regoes R. Multi-step vs. single-step resistance evolution under different drugs, pharmacokinetics, and treatment regimens. eLife. 2021;10: e64116. doi:10.7554/eLife.64116

30. Berríos-Caro E, Gifford DR, Galla T. Competition delays multi-drug resistance evolution during combination therapy. J Theor Biol. 2021;509: 110524. doi:10.1016/j.jtbi.2020.110524

31. Ojkic N, Lilja E, Direito S, Dawson A, Allen RJ, Waclaw B. A Roadblock-and-Kill Mechanism of Action Model for the DNA-Targeting Antibiotic Ciprofloxacin. Antimicrob Agents Chemother. 2020;64: e02487–19. doi:10.1128/AAC.02487-19

32. Bryson V, Szybalski W. Microbial Selection. Science. 1952;116: 45–51. doi:10.1126/science.116.3003.45

33. Timms AR, Steingrimsdottir H, Lehmann AR, Bridges BA. Mutant sequences in the rpsL gene of Escherichia coli B/r: Mechanistic implications for spontaneous and ultraviolet light mutagenesis. Mol Gen Genet MGG. 1992;232: 89–96. doi:10.1007/BF00299141

34. Alifano P, Palumbo C, Pasanisi D, Talà A. Rifampicin-resistance, rpoB polymorphism and RNA polymerase genetic engineering. J Biotechnol. 2015;202: 60–77. doi:10.1016/j.jbiotec.2014.11.024

35. Huseby DL, Pietsch F, Brandis G, Garoff L, Tegehall A, Hughes D. Mutation supply and relative fitness shape the genotypes of ciprofloxacin-resistant *Escherichia coli*. Mol Biol Evol. 2017; msx052. doi:10.1093/molbev/msx052

36. Foster PL. Methods for Determining Spontaneous Mutation Rates. In: Judith L. Campbell and PM, editor. Methods in Enzymology. Academic Press; 2006. pp. 195–213. Available: http://www.sciencedirect.com/science/article/pii/S0076687905090129

37. Wickens HJ, Pinney RJ, Mason DJ, Gant VA. Flow Cytometric Investigation of Filamentation, Membrane Patency, and Membrane Potential in *Escherichia coli* following Ciprofloxacin Exposure. Antimicrob Agents Chemother. 2000;44: 682–687. doi:10.1128/AAC.44.3.682-687.2000

38. Bos J, Zhang Q, Vyawahare S, Rogers E, Rosenberg SM, Austin RH. Emergence of antibiotic resistance from multinucleated bacterial filaments. Proc Natl Acad Sci. 2015;112: 178–183. doi:10.1073/pnas.1420702111

39. Siegel EC, Bryson V. Mutator Gene of *Escherichia coli* B. J Bacteriol. 1967;94: 38–47. doi:10.1128/jb.94.1.38-47.1967

40. Lee H, Popodi E, Tang H, Foster PL. Rate and molecular spectrum of spontaneous mutations in the bacterium *Escherichia coli* as determined by whole-genome sequencing. Proc Natl Acad Sci. 2012;109. doi:10.1073/pnas.1210309109

41. Foster PL, Lee H, Popodi E, Townes JP, Tang H. Determinants of spontaneous mutation in the bacterium *Escherichia coli* as revealed by whole-genome sequencing. Proc Natl Acad Sci. 2015;112. doi:10.1073/pnas.1512136112

42. Sun L, Alexander HK, Bogos B, Kiviet DJ, Ackermann M, Bonhoeffer S. Effective polyploidy causes phenotypic delay and influences bacterial evolvability. De Visser A, editor. PLOS Biol. 2018;16: e2004644. doi:10.1371/journal.pbio.2004644

43. Carballo-Pacheco M, Nicholson MD, Lilja EE, Allen RJ, Waclaw B. Phenotypic delay in the evolution of bacterial antibiotic resistance: Mechanistic models and their implications. Tanaka MM, editor. PLOS Comput Biol. 2020;16: e1007930. doi:10.1371/journal.pcbi.1007930

44. Hayward RS. DNA Blockade by Rifampicin-Inactivated *Escherichia coli* RNA Polymerase, and Its Amelioration by a Specific Mutation. Eur J Biochem. 1976;71: 19–24. doi:10.1111/j.1432-1033.1976.tb11084.x

45. Edgar R, Friedman N, Molshanski-Mor S, Qimron U. Reversing Bacterial Resistance to Antibiotics by Phage-Mediated Delivery of Dominant Sensitive Genes. Appl Environ Microbiol. 2012;78: 744–751. doi:10.1128/AEM.05741-11

46. Wang X, Li W, Wang W, Wang S, Xu T, Chen J, et al. Involvement of Small Colony Variant-Related Heme Biosynthesis Genes in Staphylococcus aureus Persister Formation in vitro. Front Microbiol. 2021;12: 756809. doi:10.3389/fmicb.2021.756809

47. Timms AR, Dewan KK, Bridges BA. Growth rate effects of mutations conferring streptomycindependence and of ancillary mutations in the *rpsL* gene of *Escherichia coli* : implications for the clustering (hypermutation) hypothesis for spontaneous mutation. Mutagenesis. 1995;10: 463–466. doi:10.1093/mutage/10.5.463

48. Wolff E, Kim M, Hu K, Yang H, Miller JH. Polymerases Leave Fingerprints: Analysis of the Mutational Spectrum in *Escherichia coli rpoB* To Assess the Role of Polymerase IV in Spontaneous Mutation. J Bacteriol. 2004;186: 2900–2905. doi:10.1128/JB.186.9.2900-2905.2004

49. Wu EY, Hilliker AK. Identification of Rifampicin Resistance Mutations in *Escherichia coli*, Including an Unusual Deletion Mutation. Microb Physiol. 2017;27: 356–362. doi:10.1159/000484246

50. Stracy M, Snitser O, Yelin I, Amer Y, Parizade M, Katz R, et al. Minimizing treatment-induced emergence of antibiotic resistance in bacterial infections. Science. 2022;375: 889–894. doi:10.1126/science.abg9868

51. Lloyd DP, Allen RJ. Competition for space during bacterial colonization of a surface. J R Soc Interface. 2015;12: 20150608. doi:10.1098/rsif.2015.0608

52. Thomason LC, Costantino N, Court DL. *E. coli* Genome Manipulation by P1 Transduction. Curr Protoc Mol Biol. 2007;79. doi:10.1002/0471142727.mb0117s79

53. Merlin C, McAteer S, Masters M. Tools for Characterization of *Escherichia coli* Genes of Unknown Function. J Bacteriol. 2002;184: 4573–4581. doi:10.1128/JB.184.16.4573-4581.2002

54. Garrison E, Marth G. Haplotype-based variant detection from short-read sequencing. arXiv; 2012. doi:10.48550/arXiv.1207.3907

55. Rausch T, Zichner T, Schlattl A, Stütz AM, Benes V, Korbel JO. DELLY: structural variant discovery by integrated paired-end and split-read analysis. Bioinformatics. 2012;28: i333–i339. doi:10.1093/bioinformatics/bts378

56. Delahaye C, Nicolas J. Sequencing DNA with nanopores: Troubles and biases. PLOS ONE. 2021;16: e0257521. doi:10.1371/journal.pone.0257521

57. Gillespie DT. Approximate accelerated stochastic simulation of chemically reacting systems. J Chem Phys. 2001;115: 1716–1733.

