## Supplementary Figures for "Simple theoretical models predict some - but not all – aspects of the experimental evolution of antibiotic resistance"

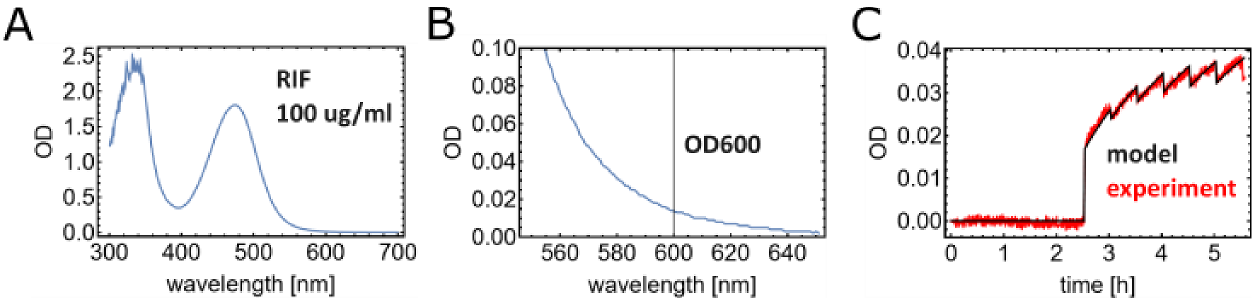


**Figure S1. Rifampicin contribution to the optical density.** (A) Absorption spectrum of freshly prepared rifampicin (100 ug/ml) in LB. (B) A range of the same spectrum magnified to show absorption at 600 nm - the wavelength used in the spectrophotometer to measure the optical density. RIF is thus expected to contribute about 0.02 to the total OD, which is about 20% of the maximum expected OD in our bioreactor experiments. (C) OD versus time for a bioreactor experiment in which rifampicin was injected at t=2.5 h and added during subsequent dilution steps to maintain the concentration of 100 ug/ml. The measured optical density changes in time due to rifampicin slowly converted to a more absorbing product (rifampicin quinone) with rate constant lambda. Fitting the model to data (3 replicates; only one shown here for clarity) gives lambda=0.5 $h^{-1}$ and the RIF-quinone contribution to the OD = 3.3 times the contribution of pure RIF. The decay rate lambda is different in the presence of bacteria (lambda=0.2 $h^{-1}$, obtained from model fitting to long-term evolutionary experiments, see Methods); this rate was used to subtract RIF contribution from OD vs time curves in Fig. 2.

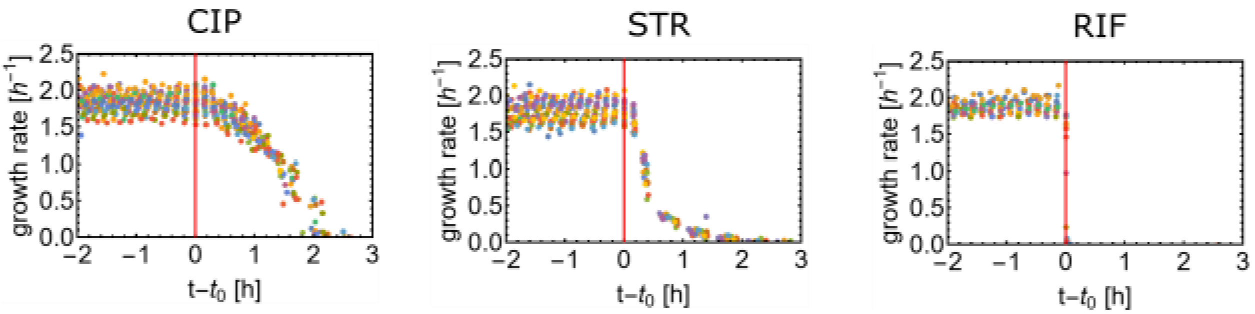


**Figure S2. Comparison of the response times for CIP, STR, and RIF**. Plots show the growth rate before and after the antibiotic (vertical red line at t=0).

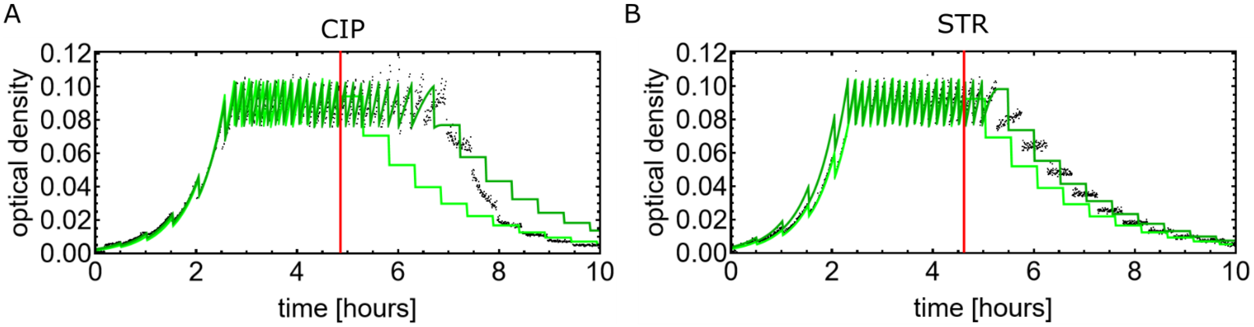


**Figure S3. The model with time-varying cell-density to optical density conversion factor, for CIP and STR.** Black = experimental curves. Light green = model with constant conversion factor (as in Fig. 2C). Dark green = model with time-dependent factor.

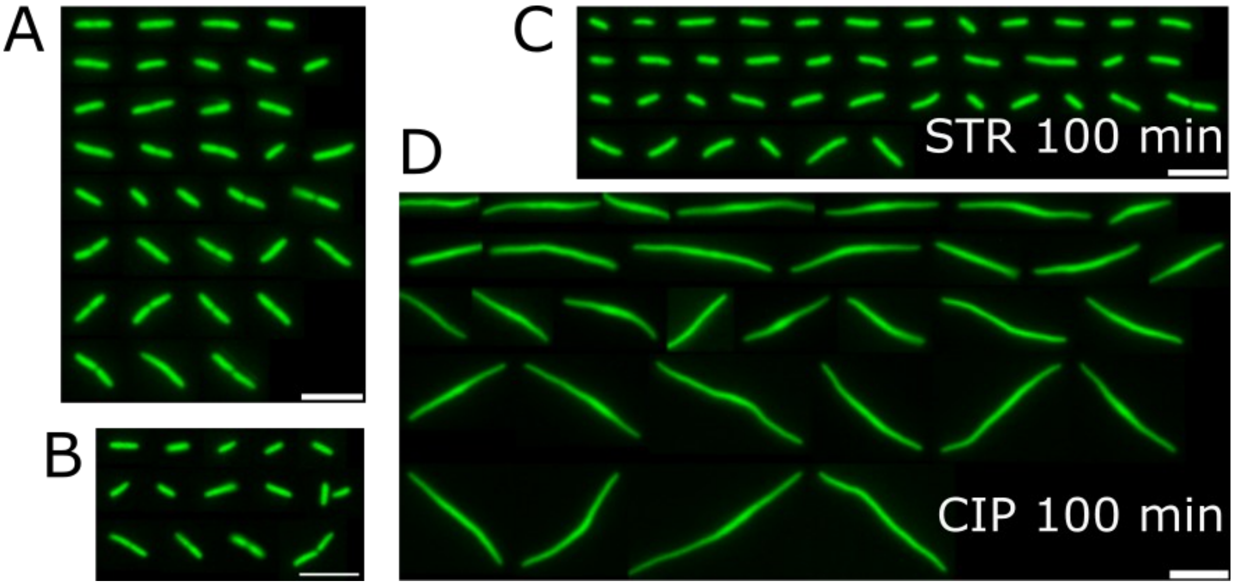


**Figure S4. Fluorescent images of individual cells before and after antibiotic exposure. (A-B)** Cells before STR (A) and CIP (B) exposure. **(C)** Cells 100 min after the exposure to STR. **(D)** Cells 100 min after the exposure to CIP. Yellow-fluorescent strain EEL02 was used in each case, and the white scale bar = 10 μm. Imaging settings: 40x NA 0.9 air objective, exposure time: 300 ms, camera: Andor Zyla 4.2+, filter: FIT-C, excitation lamp: CoolLED pE300 at 20% power. Bacteria were deposited on a glass slide and immediately covered with a cover slip #1. Images of well-focused individual cells were cropped, rotated 90° whenever required to make the cells better aligned horizontally, and assembled into a collage. Only a fraction of all imaged cells is shown for each condition.

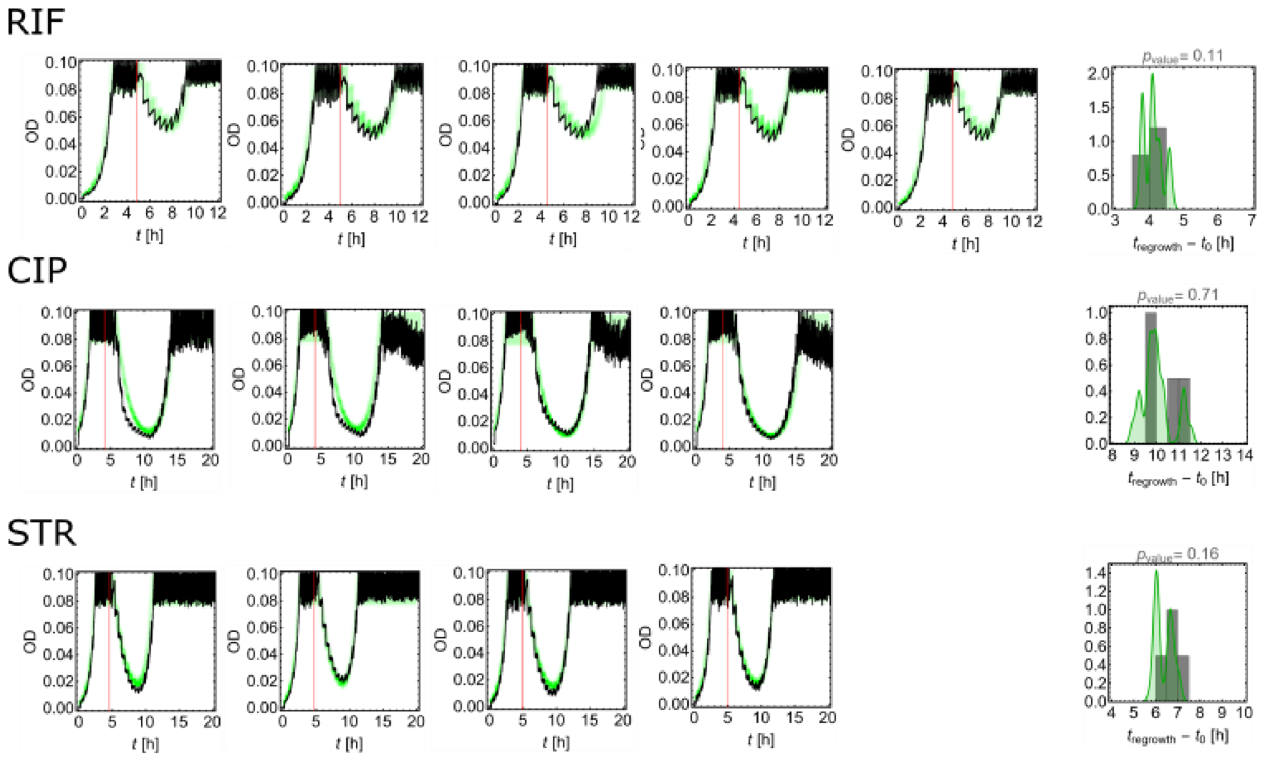


**Figure S5. The simple model correctly reproduces resistant regrowth in a standing variation experiment.** Replicates of OD versus time for bioreactor experiments in which a sensitive population spiked-in with a small number of resistant mutant cells was exposed to RIF, CIP, and STR. Black = experiment, green = model prediction (1000 simulations presented as the probability heatmap). The red line marks the beginning of antibiotic exposure. Mutants used and their initial fractions (determined by plating): RIF: *rpoB* D516Y, ratio 1:1000; CIP: *gyrA* S83L, ratio 1:1000; STR: *rpsL* K43T, ratio 1:1660. The right-most column shows regrowth time distributions and the p-values for KS tests comparing the experimental and predicted distributions.

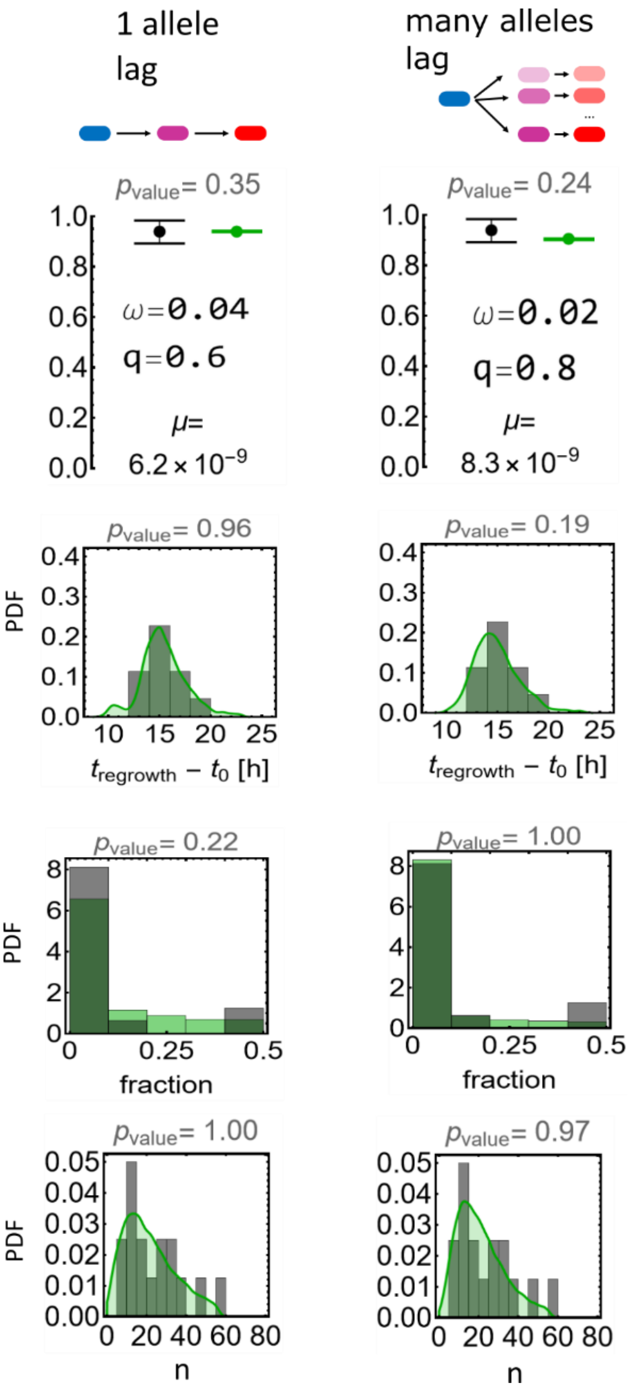


**Figure S6. Additional models for RIF.** Rows of plots show: the probability of resistant regrowth, the time to regrowth, the fraction of the least abundant strain at the end of resistant regrowth, and the distribution of the number of resistant mutants prior to antibiotic exposure. Grey and green histograms are the experimental and simulated data, respectively.

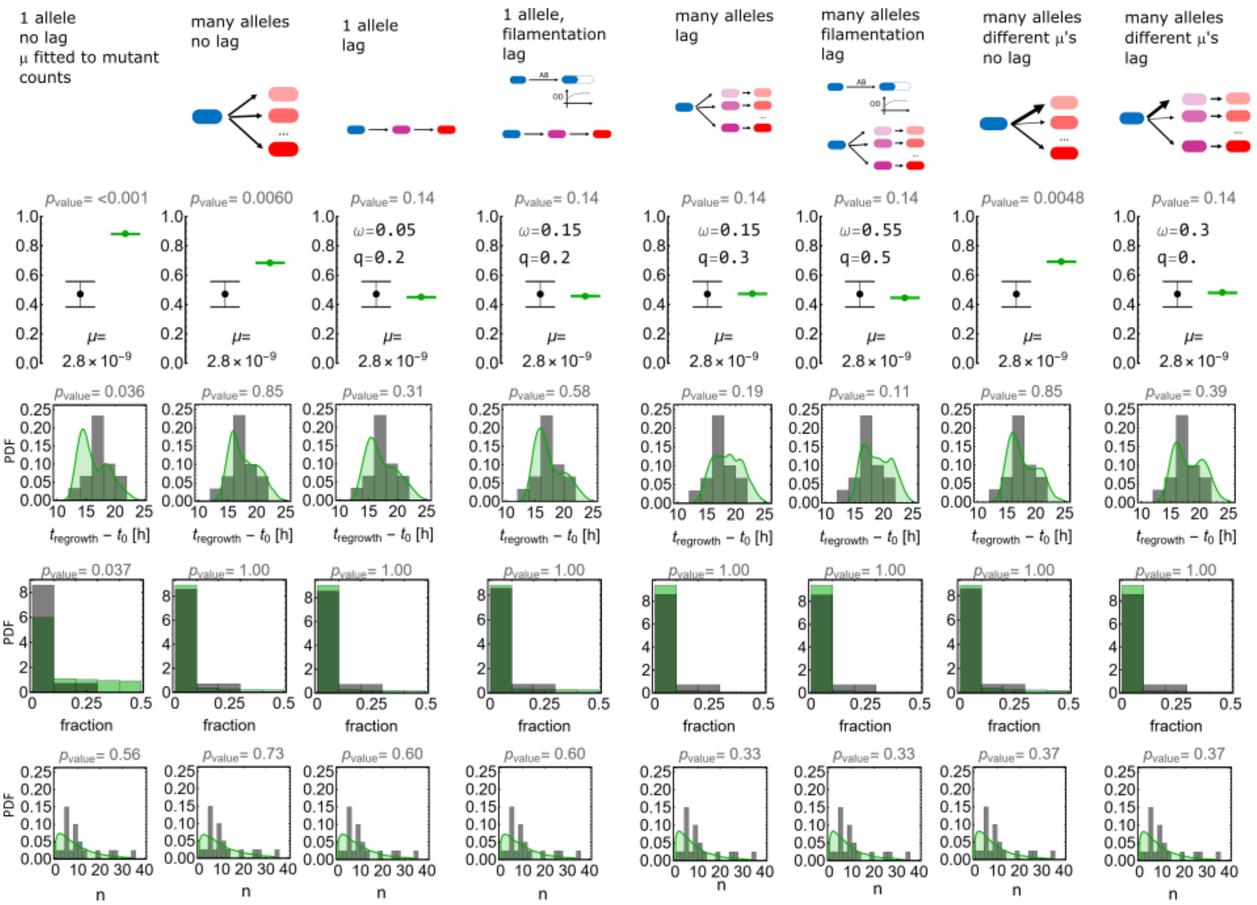

**Figure S7. Additional models for CIP.** Rows of plots show: the probability of resistant regrowth, the time to regrowth, the fraction of the least abundant strain at the end of resistant regrowth, and the distribution of the number of resistant mutants prior to antibiotic exposure. Grey and green histograms are the experimental and simulated data, respectively.
